# Tracing cellular heterogeneity in pooled genetic screens via multi-level barcoding

**DOI:** 10.1101/224345

**Authors:** Michael Boettcher, Sergio Covarrubias, Anne Biton, James A Blau, Haopeng Wang, Noah Zaitlen, Michael T McManus

## Abstract

While pooled loss- and gain-of-function screening approaches have become increasingly popular to systematically investigate mammalian gene function, they have thus far ignored the fact that cell populations are heterogeneous. Here we introduce multi-level barcoded sgRNA libraries to (i) monitor differences in the behavior of multiplexed clonal cell lines, (ii) trace sub-clonal lineages of cells expressing the same sgRNA, (iii) derive in-sample screen replicates and (iv) reduce the number of cells and sequencing read counts required to reach statistical significance. Using our approach, we illustrate how clonal heterogeneity impairs the results of pooled genetic screens and demonstrate the ability of multi-level barcoding to resolve cellular heterogeneity related issues.

## Background

Pooled genetic screens are a powerful tool to functionally dissect genetic networks in mammalian cells and in conjunction with recently developed CRISPR/Cas systems, they permit a variety of scalable genetic perturbations, including gene knockout, knockdown or activation [1]. While numerous pooled CRISPR screens have been conducted successfully in the past, they frequently disregard a fundamental property of cell populations – namely their genotypic and phenotypic heterogeneity [2]. As a matter of fact, most CRISPR screens published to date were conducted in clonal Cas9 lines derived from single cells [1, 3–7], thereby restricting cellular heterogeneity and biasing screen results.

To systematically dissect the influence of cellular heterogeneity on pooled genetic screens, we developed a three-level barcoding strategy for pooled sgRNA libraries. Our strategy is based on the combination of every sgRNA sequence (level 1) in the library with two extra features; a random barcode nucleotide sequence (BC, level 2) and a constant library identifier (ID, level 3). The ID consists of a sequence that is unique to each sgRNA library within which each sgRNA is associated with one of ~100 randomly generated BC sequences. BC sequences hence allow the analysis of sub-clonal lineages of cells expressing a certain sgRNA sequence, similar to recently described random sequence labels [8] or unique molecular identifiers [9]. Beyond that; the library specific ID facilitates the tracing of multiplexed clonal Cas9 lines over the course of pooled screens and thereby the performance of clonal screen replicates under identical conditions.

Here, we describe the utility of multi-level barcoded sgRNA libraries for pooled genome-scale CRISPR-mediated gene knockout (CRISPRwt) as well as knockdown (CRISPRi) screens to systematically identify genes involved in TRAIL-mediated apoptosis [10]. Using our barcoding approach, we demonstrate that (i) clonal Cas9 lines show heterogeneous responses to TRAIL-receptor (TRAIL-R) antibody treatment under identical experimental conditions and (ii) clonal variations in treatment resistance induce substantial differences in statistical power and thereby detectable candidate genes. Similar issues caused by clonal heterogeneity likely bias the outcomes of many if not all genetic screens. Here we demonstrate the utility of multi-level barcoding to overcome limitations of conventional screening approaches by dissecting cellular heterogeneity in multiplexed genetic screens.

## Results

Either Cas9 nuclease (CRISPRwt) or dCas9-KRAB (CRISPRi) was introduced into a population of Jurkat T lymphocytes. Clonal lines from either system were expanded and the function of their respective CRISPR systems was confirmed as previously described [11]. Although all clonal lines were derived from the same parental population, they displayed substantial variability in their response to TRAIL-R (TNFRSF10B) antibody treatment (Figure 1A). To investigate the impact of the observed heterogeneity on CRISPR screens, one resistant (CloneR) and one sensitive clone (CloneS) from either CRISPR system was used for pooled screens to identify genes involved in TRAIL-mediated apoptosis (Figure 1B). Each clonal line was transduced with one of four barcoded sgRNA libraries, targeting every protein coding gene in the human genome for knockout (CRISPRwt) or knockdown (CRISPRi). To enable tracing of the relative abundance of the clonal Cas9 lines, each sgRNA library was tagged with one of four different IDs. After lentiviral transduction of the sgRNA libraries at low MOI, followed by the selection of successfully transduced cells, the four clonal lines were multiplexed at equal numbers, ensuring a representation of each sgRNA in over 1,000 cells from each clonal Cas9 line. The cells were then split into two bioreactor-vessels: one untreated control vessel and one vessel treated with escalating doses of TRAIL-R antibody on days 0, 2 and 4. Cells from the beginning of the screen (baseline) as well as from days 4, 9 and 14 were harvested from both vessels. ID, BC and sgRNA sequences from each time point were recovered via PCR and quantified by means of paired-end next generation sequencing (Figure 1C).

**Figure 1.**
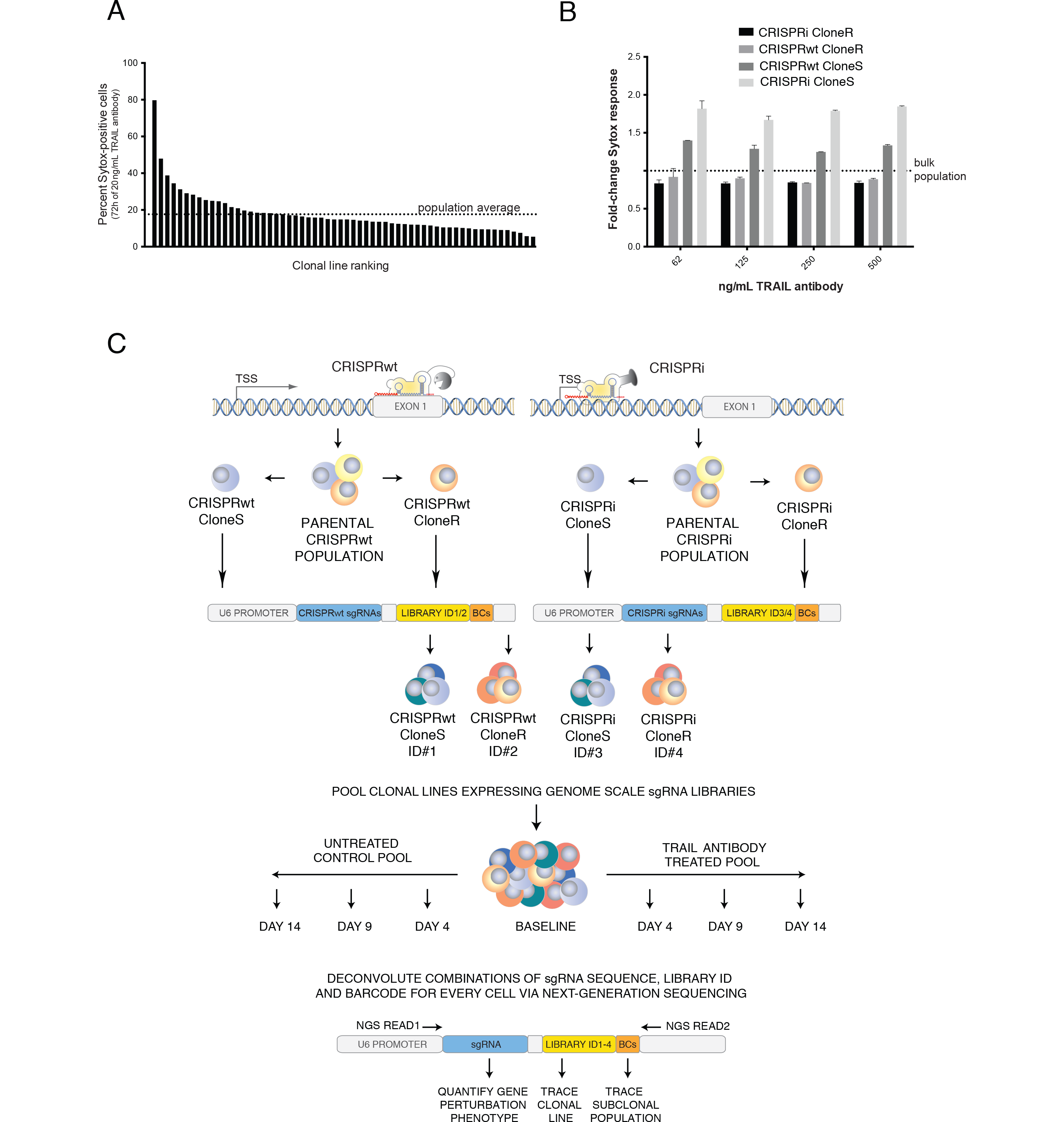
**A**. Apoptosis response of 60 clonal Cas9 Jurkat cell lines following 72h treatment with 20 ng/μL TRAIL-R antibody was assessed by sytox staining and FACS analysis. **B**. Apoptosis response of the four clonal Cas9 lines used for pooled CRISPR screens at indicated concentrations of TRAIL-R antibody. **C**. Schematic of multiplexed CRISPR screening and deconvolution approach. For CRISPRwt (top left) and CRISPRi (top right) the respective Cas9 system was introduced into a population of Jurkat cells followed by the characterization of clonal lines for functionality of each CRISPR system. From both systems, two clonal lines were transduced with a multi-level barcoded sgRNA library (12 sgRNAs/gene) to knockout (CRISPRwt) or knockdown (CRISPRi) each protein coding gene in the human genome. Successfully transduced cells were multiplexed at equal numbers and the abundance of each clonal Cas9 line was traced via one of four library identifier sequences (ID) throughout the screen. Cell pools were cultured for 14 days in the absence (bottom left) or presence (bottom right) of TRAIL-R antibody. For downstream analysis via next-generation sequencing, sgRNA expression cassettes including the sgRNA encoding sequence, ID and BCs were recovered via PCR from the genomic DNA of cell pools from the beginning of the screen (baseline) as wells as on days 4, 9 and 14. Using a paired-end sequencing strategy allowed the quantification of the dis-/enrichment of sub-clonal populations (BC) within clonal lines (library ID) following the perturbation of any protein-coding gene in the human genome (sgRNA sequence).

Utilizing the unique sgRNA library IDs, enrichment of both resistant clones was observed over time when compared to both sensitive clones after TRAIL-R antibody treatment (Figure 2A and Supplementary Table S1). Those results confirm the aforementioned heterogeneous apoptosis response of the four clonal Cas9 lines (Figure 1B) and illustrate the suitability of ID sequences to trace clonal lines in pooled cell populations. In order to identify genes involved in TRAIL-mediated apoptosis, all four CRISPR screens were analyzed using MAGeCK [12]. A higher number of significantly enriched candidate genes (FDR < 5%) was detected in the CRISPRwt compared to CRISPRi screens and in ClonesS compared to ClonesR (Figure 2B, Supplementary Tables S2-5). These results are consistent with the claim that CRISPRwt outperforms CRISPRi in identifying essential genes [7].

**Figure 2.**
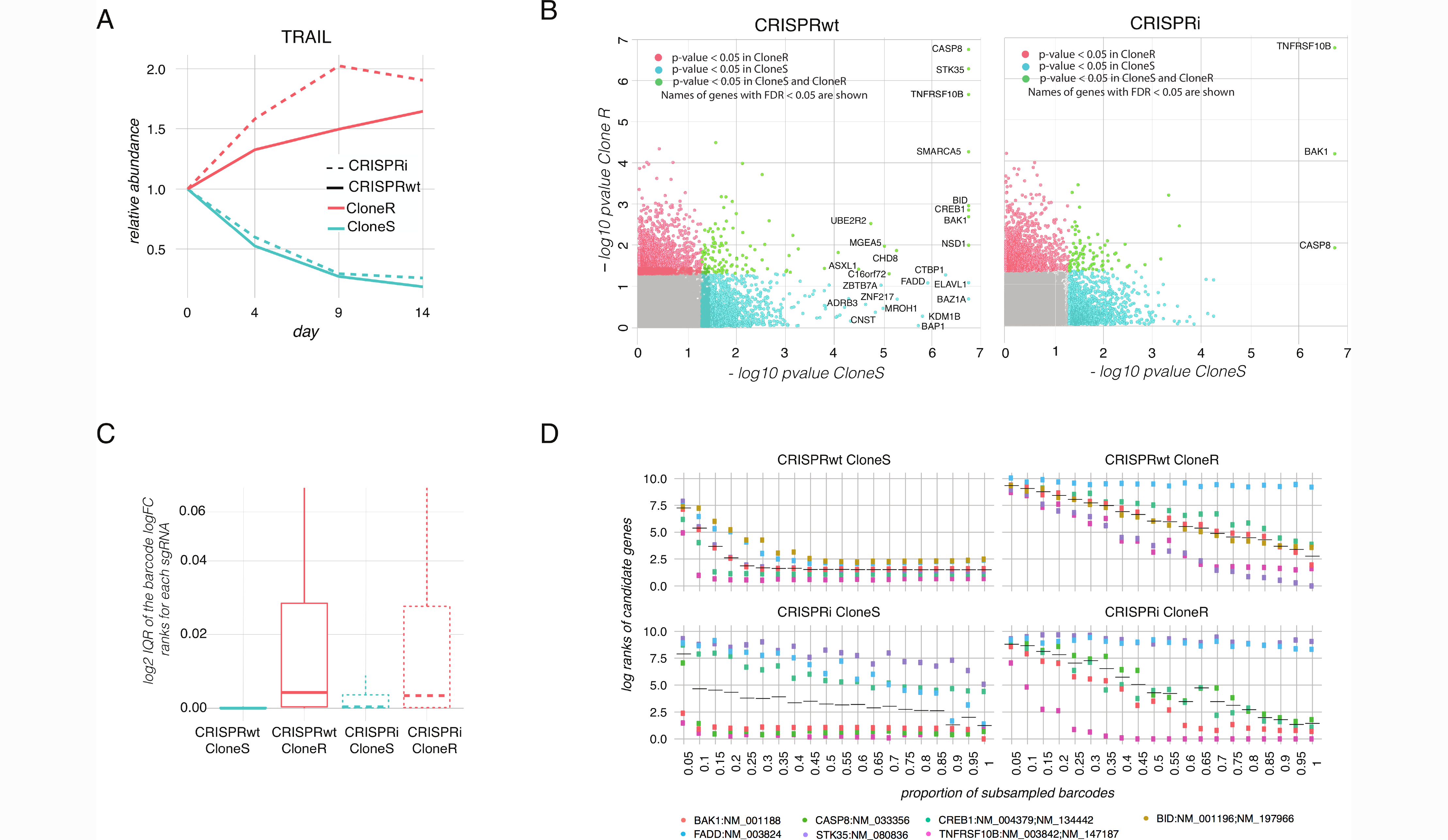
**A.** Relative abundance of clonal CRISPRwt and CRISPRi populations in the TRAIL-R antibody treated cell pool. Abundance of library IDs normalized to the baseline (day 0) is shown on the y-axis. **B**. Summary of multiplexed screen results. The -log_10_ p-values obtained from MAGeCK analysis of TRAIL-R antibody treated versus untreated cell pools on day 14 are shown. Genes highlighted in green have p-values below 0.05 in cloneR and cloneS. Genes with p-values below 0.05 only in cloneS are shown in blue. Genes with p-values below 0.05 only in cloneR are shown in red. Names of genes with a FDR < 5% are indicated. **C**. The distribution of BC log fold changes for each sgRNA is shown for each of the four clonal Cas9 lines. Interquartile ranges (IQR) of the ranks of log fold-changes for each sgRNA across barcodes are shown. **D**. Barcode subsampling across the four clonal Cas9 lines. The x-axis shows the fraction of barcodes sampled for each sgRNA. The y-axis shows the log_10_ rank of candidate genes in the MAGeCK output obtained from the subsampled datasets.

Next, we explored the utility of random barcodes (BCs) associated with each sgRNA sequence to (i) derive screen replicates, (ii) trace individual sub-clonal populations and (iii) simulate the impact of reduced screen complexity on screen results. First, all BCs were randomly split into three bins, providing three in-sample replicates. High levels of correlation between in-sample replicates confirm that the randomly assembled sub-populations within each clonal Cas9 line showed an overall similar behavior (Supplementary Figure 1). Next, all individual sgRNA-BC sequence combinations were analyzed separately to further resolve the influence of sub-clonal heterogeneity on the outcomes of CRISPR screens. Figure 2C shows a summary of the variation within individual sub-populations across the four clonal Cas9 lines. The distribution of all BC fold-changes associated with a given sgRNA is shown as interquartile ranges (IQRs). Overall, the resistant clones showed a wider distribution of BC fold-changes when compared to the sensitive clones. These results suggest the existence of larger sub-clonal variation in response to TRAIL-R antibody treatment in resistant compared to sensitive clones.

Two major concerns for every pooled CRISPR screen are (i) sgRNA library representation in the target cells and (ii) sgRNA sequence read depth. For screens conducted with large sgRNA libraries or in difficult to culture cells, these two issues can rapidly become limiting factors and in some scenarios, such as *in-vivo* screens, it is even impossible to determine sgRNA library representation. Hence assessing and reducing the complexity of genetic screens, without compromising statistical power is critical to success in many challenging screen setups. To investigate the effects of reduced complexity on the screen results, we sub-sampled fractions of BCs associated with each sgRNA in the library and determined the minimum fraction required to significantly call sgRNAs identified from the full read counts. As shown in Figure 2D on the example of the candidate genes TNFRSF10B (TRAIL receptor), BAK1, BID, CASP8, CREB1, FADD and STK35, only 25% of sequencing reads were sufficient to identify all candidate genes in CloneS of the CRISPRwt screen while in CloneR over 60% of the reads were necessary to call even the strongest hits TNFRSF10B, CASP8 and STK35. For FADD, even the inclusion of 100% of the BCs did not result in a significant rank (Figure 2D). Similar results were obtained from both clones of the CRISPRi screens. These findings illustrate how apoptosis-resistant clonal Cas9 lines, because of their aforementioned heterogeneity, require higher screen complexity to capture an enrichment signal similar to the one of sensitive clonal Cas9 lines. More importantly, these results confirm the utility of BCs to determine the level of saturation in genetic screens.

## Conclusion

Here we describe a multi-level barcoding system to readily trace clonal cell lines as well as their sub-clonal populations in pooled genetic screens. Our approach expands on recently published methods for capturing cellular heterogeneity via sub-clonal CRISPR screen analysis [8, 9] by additionally introducing sgRNA library IDs to trace multiplexed clonal Cas9 cell lines under identical screen conditions. We demonstrate how this combinatorial barcoding approach addresses experimental variability caused by clonal heterogeneity and thereby increases statistical power to detect candidate genes, ultimately reducing the amount of cells and sequencing read counts to reach statistical significance. Here we exemplify the utility of multi-level barcoding by means of CRISPR knockout and CRISPRi screens; yet our approach is applicable to resolve issues with cellular heterogeneity in virtually any type of pooled genetic screen including RNAi, cDNA or CRISPRa screens. The pipelines and resources developed here, including multi-level barcoded sgRNA expression vector systems and genome-scale CRISPRwt/i libraries will allow researchers to analyze future genetic screens at clonal and sub-clonal levels, thereby adding an unprecedented layer of control to pooled genetic screens analysis.

## Methods

### Clonal Jurkat cell lines

Clonal Jurkat CRISPRwt and CRISPRi lines were derived as previously described [11] and cultured in RPMI 1640 medium, supplemented with 10% fetal bovine serum and 1x Anti-Anti (Gibco).

### Sytox apoptosis assay

Jukat cells were treated with TRAIL receptor antibody (MAB631, R&D Systems) for indicated periods of time. Cells were then stained with a 1:500 dilution of Sytox Green (Thermofisher, S7020) and analyzed on a flow cytometer to quantify the fraction of GFP+ (dead) cells.

### CRISPRwt and CRISPRi sgRNA library design

For both CRISPR technologies, a separate genome-scale sgRNA library was designed, each consisting of over 250,000 total sgRNAs targeting every unique Refseq annotated (hg19) protein coding isoform with up to 12 sgRNAs, plus 7,700 non-target control sequences (NTC). For the CRISPRwt sgRNA library, the earliest possible exon of each transcript variant was targeted. For the CRISPRi sgRNA library, sgRNAs were targeted 50 to 500 bp downstream of the transcription start site (TSS) of each isoform. All sgRNAs were designed against target sites that are of the format (N)_20_NGG, and selected sgRNAs must pass the following off-targeting criteria: 1) the 11bp-seed must not have an exact match in any other promoter region, and 2) if there is an exact off-target seed match, then the rest of the sgRNA must have at least 7 mismatches with the potential off-target site. After all sgRNAs that pass off-targeting criteria were generated, up to 12 sgRNAs/transcript were selected. All sgRNA sequences are shown in Supplementary Tables S6 (CRISPRwt) and S7 (CRISPRi).

In addition to the sgRNA sequence, every library plasmid contained two extra features that allowed us to address heritable clonal heterogeneity in CRISPR screens: 1.) A specific 6 nucleotide long library identifier (ID) sequence (IDs for CRISPRwt libraries were ‘GCCTAA’ or ‘TGGTCA’, and for CRISPRi libraries ‘CGTGAT’ or ‘ACATCG’ respectively) to allow tracing of clonal lines in a pool cells. And 2.) a unique 20 nucleotide barcode sequence to facilitate the analysis of sub-clonal populations (see Extended Data 1 for vector map).

### sgRNA library cloning

For both, the CRISPRwt and CRISPRi libraries, the designed 20 nt target specific sgRNA sequences were synthesised as a pool, on microarray surfaces (CustomArray, Inc.), flanked by overhangs compatible with Gibson Assembly into the pSico based barcoded sgLenti sgRNA library vector (see Extended Data 1 for vector map). The synthesised sgRNA template sequences were of the format: 5’-GGAGAACCACCTTGTTGG-(N)_20_-GTTTAAGAGCTATGCTGGAAAC-3’. Template pools were PCR amplified using Phusion Flash High-Fidelity PCR Master Mix (ThermoFisher Scientific) according to the manufacturers protocol with 1 ng/μL sgRNA template DNA, 1 μM forward primer (5’-GGAGAACCACCTTGTTGG-3’), 1 μM reverse primer (5’-GTTTCCAGCATAGCTCTTAAAC-3’) and the following cycle numbers: 1x (98C for 3 min), 15x (98C for 1 sec, 55C for 15 sec, 72C for 20 sec) and 1x (72C for 5 min). PCR products were purified using Minelute columns (Qiagen). The library vector sgLenti was prepared by restriction digest with AarI (Thermo-Fischer) at 37C overnight, followed by 1% agarose gel excision of the digested band and purification via NucleoSpin columns (Macherey-Nagel). Using Gibson Assmbly Master Mix (NEB), 1000 ng digested sgLenti and 100 ng amplified sgRNA library insert were assembled in a total 200 μL reaction volume. The reaction was purified using P-30 buffer exchange columns (Biorad) that were equilibrated 5x with H_2_O and the total eluted volume was transformed into three vials of Electromax DH5α (ThermoFisher). E.coli were recovered, cultured overnight in 500 mL LB (100 ug/mL ampicillin) and used for Maxiprep (Qiagen). In parallel, a fraction of the transformation reaction was plated and used to determine the total number of transformed clones. The library cloning coverage (number of E.coli colonies per sgRNA plasmid) was determined to be >100x for each of the four libraries, ensuring even representation of all library sgRNA sequences and their narrow distribution (Extended Data Fig. QC) as well as the required barcode diversity for each sgRNA sequence to facilitate the tracing of sub-clonal populations.

### Lentivirus production

HEK293T cells were seeded at 65,000 cells per cm^2^ in 15 cm dishes in 20 mL medium (DMEM, 10% fetal bovine serum) and incubated overnight at 37C, 5% CO_2_. The next morning, 8 ug sgRNA library plasmid, 4 ug psPAX2 (Addgene #12260), 4 ug pMD2.G (Addgene #12259) and 40 μL jetPRIME (Polyplus) were mixed into 1 mL serum free OptiMEM (Gibco) with 1x jetPRIME buffer, vortexed and incubated for 10 min at RT and added to the cells. 24 h later, 40U DNAseI (NEB) were added to each plate in order to remove untransfected plasmid and at 72h post-transfection, supernatant was harvested, passed through 0.45 μm filters (Millipore, Stericup) and aliquots were stored at -80C.

### Genome-wide CRISPRwt/CRISPRi screens

Two clonal CRISPRwt and two CRISPRi Jurkat lines were transduced separately with their respective sgRNA libraries (ID 1-4) at low multiplicity of infection (MOI=0.3) to reduce the frequency of multiple-infected cells; thus, only one gene was targeted for knockout or knockdown in each cell. The library coverage at transduction was determined to be ~100 transduced cells for each sgRNA from each of the four libraries, to ensure full representation of library sgRNA sequences in the target cell populations as well as a sufficient diversity of barcode sequences per sgRNA to facilitate sub-clonal population analyses. The four transduced clonal lines were then cultured separately in RPMI with 10% FBS and 1x Anti-Anti (Gibco) in a 37°C incubator with 5%CO_2_. 2 days post transduction, cells were selected for 5 days with puromycin (2 ug/mL). Following puromycin selection, 1 billion cells from each of the four clonal CRISPRwt/i lines were multiplexed. The cell pool was seeded into 2x 5 litre RPMI medium in CelliGen BLU bioreactor vessels (25 RPM, 37°C, pH=7.4, O_2_=8%) at a final density of 300,000 cells per ml - or a total of 1.5 billion cells per bioreactor - to achieve a representation of >1,000 cells per sgRNA in each of the four libraries. The remaining pool of 1 billion cells was cryo-preserved in 90% FBS, 10% DMSO for later analyses (Baseline). After pooling, cells were either treated with escalating concentrations of TRAIL receptor antibody (MAB631, R&D Systems) on day 0 (10 ng/ml), day 1 (20 ng/ml) and day 4 (25 ng/ml) or left untreated (control cells). The culture was diluted with fresh medium when exceeding a density of 1 million cells/ml medium and a library coverage of >1,000x was maintained throughout the screen to ensure equal representation of sgRNAs and barcodes. Identical to the baseline sample, aliquots of 1 billion cells from TRAIL-R antibody treated and untreated bioreactors were cryo-preserved on days 4, 9 and 14 for later analysis.

### Genomic DNA (gDNA) extraction

Cell pellets from baseline and TRAIL-R antibody treated samples were resuspended in 20 mL P1 buffer (Qiagen) with 100 ug/mL RNase A and 0.5% SDS followed by incubation at 37C for 30 min. After that, Proteinase K was added (100 ug/mL final) followed by incubation at 55C for 30 min. After digest, samples were homogenised by passing them three times through a 18G needle followed by three times through a 22G needle. Homogenised samples were mixed with 20 mL Phenol:Chlorophorm:Isoamyl Alcohol (Invitrogen #15593-031), transferred into 50 mL MaXtract tubes (Qiagen) and thoroughly mixed. Samples were then centrifuged at 1,500g for 5 min at room temperature (RT). The aqueous phase was transferred into ultracentrifuge tubes and thoroughly mixed with 2 mL 3M sodium acetate plus 16 mL isopropanol at RT before centrifugation at 15,000g for 15 min. The gDNA pellets were carefully washed with 10 mL 70% ethanol and dried at 37C. Dry pellets were resuspended in H_2_O and gDNA concentration was adjusted to 1 ug/uL. The degree of gDNA shearing was assessed on a 1% agarose gel and gDNA was sheared further by boiling at 95C until average size was between 10-20 kb.

### PCR recovery of sgRNA sequences from gDNA

Multiple PCR reactions were prepared to allow amplification of the total harvested gDNA from a 1000x cell coverage for each sample. For the first round of two nested PCRs, the total volume was 100 μL containing 50 ug sheared gDNA, 0.3 μM forward (5’-ggcttggatttctataacttcgtatagca-3) and reverse (5’-cggggactgtgggcgatgtg-3’) primer, 200 μM each dNTP, 1x Titanium Taq buffer and 1 μL Titanium Taq (Clontech). PCR cycles were: 1x (94C - 3 min), 16x (94C - 30 sec, 65C – 10 sec, 72C – 20 sec), 1x (68C – 2 min). All first round PCRs were pooled and a fraction was used as template for the second round PCR. The total volume of the second round PCR was 100 μL containing 2 μL pooled first round PCR, 0.5 μM forward (5’-AATGATACGGCGACCACCGAGATCCACAAAAGGAAACTCACCCTAAC-3’) and reverse (5’-CAAGCAGAAGACGGCATACGAGAT-(N)6-GTGACTGGAGTTCAGACGTG-3’) primer where (N)_6_ is a 6 nt index for sequencing on the Illumina HiSeq platform, 200 μM each dNTP, 1x Titanium Taq buffer and 1 μL Titanium Taq (Clontech). PCR cycles were: 1x (94C - 3 min), 16x (94C - 30 sec, 55C – 10 sec, 72C – 20 sec), 1x (68C – 2 min). The resulting PCR product (344 bp) was extracted from a 1% agarose gel. Gel extracted bands were submitted for sequencing on an Illumina HiSeq 2500 platform using paired end 50 kits with the custom sequencing primer 5’-GAGACTATAAGTATCCCTTGGAGAACCACCTTGTTGG-3’ for reading the sgRNA sequence and the standard Truseq Illumina reverse primer to read out 20 nt unique barcode sequences and library IDs 1-4.

### Sequencing reads preprocessing

The sgRNA, library IDs and BC sequence information obtained through paired-end next generation sequencing was extracted from every read sequence using a python script (https://github.com/quasiben/gRNA Tool). Reads with the same combination of sgRNA sequence, library IDs, and BC sequence were summed up together to generate a BC-based count matrix for each library-ID.

### Barcode built-in replicates and analysis of the CRIPSR screens

In order to keep sgRNAs with a number of read counts sufficient to be used for built-in replication, we discarded the sgRNAs with less than an average of 100 read counts (summed up across barcodes) in the untreated samples from days 9 and 14. In addition only sgRNAs for which at least 3 barcodes were detected were included in the analysis. This led to the selection of 177,524 and 228,321 sgRNAs targeting 32,446 and 37,147 RefSeq gene ids for CRISPRwt in cloneS and cloneR, respectively; and 161,932 and 186,816 sgRNAs targeting 18,664 and 18,705 RefSeq ids for CRISPRi in CloneS and CloneR respectively. Barcodes were randomly split into three groups for each sgRNA. Read counts of the barcodes belonging to a same group were summed up and used as one built-in replicate. Only sgRNAs with at least three barcodes were included in the analysis.

The three built-in replicates were used as input for MAGeCK v0.5.5, with default parameters to detect sgRNAs that were positively enriched in the TRAIL-treated samples for each clonal population, each CRISPR system, and each time point independently. The analysis output from all time points is shown in Supplementary Table 2 (CRISPRwt, cloneR), Table 3 (CRISPRi, cloneS), Table 4 (CRISPRi, cloneR) and Table 5 (CRISPRwt, cloneS).

### Barcode subsampling to explore cell coverage

For each selected sgRNA with at least four barcodes detected at baseline, we randomly sub-sampled different proportions of the barcodes (from 5% to 100% of the barcodes, increasing by steps of 5%) and performed the MAGeCK analysis on each of the sets of sub-sampled data. Barcodes were summed up for each sgRNAs and days 9 and 14 were used as replicates since built-in replicates could not be used because of small number of available barcodes that could be obtained when small proportions of subsampling were used. Only sgRNAs with at least four barcodes were used and, when needed, proportions were rounded to the closest proportion that could be obtained with the minimum number of barcodes. Each subsampling was iterated 50 times, and the obtained gene ranks were averaged across the 50 iterations for each subsampling proportion.

## Acknowledgements

Special thanks go to members of the McManus Lab who provided critical feedback during the course of this project.

## Authors’ Contributions

The project was conceived and directed by M.B. and M.T.M. Clonal Cas9 Jurkat cell lines were created and characterized by H.W. All sytox experiments were performed by S.C. All genome-wide sgRNA libraries were designed by J.A.B. with guidance from M.B. and cloned by M.B. NGS sequencing strategy for multi-level barcoding read-out was developed by M.B. Genome-wide CRISPR screen optimisation was performed by M.B. and S.C. All screens were performed by M.B. with assistance from S.C. The computational pipelines for data analysis of screens were developed by A.B. with assistance from M.B. and N.Z. The manuscript was written by M.B. and A.B. with critical input from S.C., N.Z and M.T.M. All authors read and approved the final manuscript.

## Ethics Approval And Consent To Participate

Not applicable in this study.

## Data Availability

All relevant datasets have been made available in the supplement of this publication.

## Supplementary Figure 1

Scatter plots of sgRNA normalized log_2_ read counts, at baseline (day 0), between three built-in replicates made by randomly splitting barcodes into three groups for each sgRNA. Spearman rank correlation values between replicates are indicated.

## Table Legends

**Supplementary Table S1**. Number of read counts mapped to each clonal population within each CRISPR system.

**Supplementary Tables S2-S5**. MAGeCK outputs at the gene-level from CRISPRwt CloneR (Supplementary Table S2), CRISPRi CloneS (Supplementary Table S3), CRISPRi CloneR (Supplementary Table S4) and CRISPRwt CloneS (Supplementary Table S5). Columns are described at https://sourceforge.net/p/mageck/wiki/output/#gene summary txt.

**Supplementary Table S6**. CRISPRwt sgRNA library sequences.

**Supplementary Table S7**. CRISPRi sgRNA library sequences.

